# Broadening the ecology of fear: non-lethal effects arise from diverse responses to predation and parasitism

**DOI:** 10.1101/766477

**Authors:** D.R. Daversa, R.F. Hechinger, E. Madin, A. Fenton, A.I. Dell, E. Ritchie, J. Rohr, V.H.W. Rudolf

## Abstract

The ecology of fear demonstrates how prey responses to avoid predation cause non-lethal effects at all ecological scales. Parasites also elicit defensive responses in hosts with associated non-lethal effects, which raises the longstanding, yet unresolved question of how non-lethal effects of parasites compare with those of predators. We developed a framework for systematically answering this question for all types of predator and parasite systems. Our framework predicts that trait responses and their non-lethal effects should be strongest from predators and parasites that do not kill individuals to feed on them, but which nevertheless damage fitness. Analysing trait response data on amphibians, which have been well-studied for this area of research, showed that individuals generally responded more directly to short-term predation risks than to parasitism. Apart from studies using amphibians, there have been few direct comparisons of responses to predation and parasitism, and none have incorporated responses to micropredators, parasitoids, or parasitic castrators, or examined their long-term consequences. Addressing these and other data gaps highlighted by our general framework can advance the field toward understanding how non-lethal effects shape real food webs, which contain multiple predator and parasite species.

## Introduction

Few people watch the movie “Jaws” without experiencing a fear that can make them think twice before swimming in the ocean. Wildlife are no different. Animals avoid foraging where predators are or have been, reduce activity levels, seek refuges, and exhibit other responses in ‘fear’ of being eaten by predators [1,2]. Fear responses exemplify non-lethal effects of predators on prey traits, and are more generally called ‘trait responses’. Trait responses can trigger a broad range of additional non-lethal effects, including ‘trait-mediated effects’ that range from reduced individual fitness to trophic cascades [3–5] that can destabilize communities [6]. Wolves, for example, frighten elk away from exposed foraging grounds into sheltered habitats with less nutritious vegetation, which then reduces elk birth rates [7] and alters vegetation structure [8]. By eliciting trait responses, predators impact prey, and wider communities, without direct killing.

Recent surges in disease outbreaks make clear that parasites also elicit trait responses with non-lethal effects. To reduce infection risk, hosts may avoid infected conspecifics [9–11], defend against infectious propagule attack [12], or avoid risky areas, such as faeces representing a hot spot of undetectable nematode eggs [13–15]. Basic emotions like ‘disgust’ [15–18] and the age-old cliché “avoid like the plague” suggest that parasite avoidance is interwoven in our own history as much as is our fear of predators.

Trait responses to parasites are not confined to avoidance tactics like those made in fear or disgust. Because parasitism is not immediately lethal, hosts can respond, and incur non-lethal effects, after a successful attack by a parasite [19–21]. Immune responses are one of myriad host responses made after infection that can have non-lethal effects [22]. Post-attack trait responses elicited by parasites has led some to hypothesize that parasites cause stronger cumulative non-lethal effects than predators [5,23]. A formal framework for comparing the full range of trait responses to predators and parasites is lacking, however, leaving uncertainty over the similarities and differences in how non-lethal effects accumulate across different predator-prey and host-parasite interactions.

Here, we compare trait responses to predation and parasitism, considering how variation in their frequency and strength may drive differences in how non-lethal effects accrue in prey and hosts. Building on recent conceptual developments [21,24], our goal is to establish a quantitative foundation for estimating non-lethal effects in real ecosystems, which contain multiple predator and parasite species. We draw from consumer-resource theory to construct a general framework for estimating trait responses that accounts for the variable consumer strategies of predators and parasites. Unlike current conceptual frameworks for fear and disgust, we deconstruct interactions into sequential phases to consider trait responses before, during, and after an attack, thus allowing us to compare and contrast the complete diversity of predatory and parasitic consumers. We use this framework to form specific predictions regarding how trait responses, and their non-lethal effects, should manifest from interactions with different types of predators and parasites. For brevity, we focus on adaptive trait responses for defence against predation or parasitism, but our framework can also consider maladaptive responses, such as those occurring from parasite manipulation [25]. We then report on a systematic review of the comparative literature on trait responses. We also analysed data on larval amphibians, the most common animal group used by the reviewed studies, to assess the empirical evidence for our predictions. We conclude by highlighting unresolved questions about trait responses and their non-lethal effects on communities and ecosystems.

### A general trait-response framework for examining non-lethal effects

#### Existing trait response frameworks

The ecology of fear in predator-prey systems provides a strong, yet incomplete foundation for examining trait responses and their impact on population and community dynamics. ‘Fear responses’ denote trait responses to the risk of predation, before a predator attack. Prey movement away from foraging habitats when predators are nearby is a well-studied ‘fear response’ [1,3,26]. As Lima and Dill (1990) pointed out in their seminal framework [1], prey may also exhibit defensive trait responses during predator attack and even after being captured, phases not covered within the standard domain of the ‘ecology of fear’. Systematic examination of non-lethal effects must go beyond fear to consider multiple trait responses made throughout interactions. This becomes especially obvious when incorporating responses to parasites.

In stark contrast to predation, host responses after parasite attack, that is, after infection, can continue as parasites feed on individuals. The same is true for prey of micropredators, which act like parasites by feeding without killing. This distinguishing characteristic of host-parasite and micropredator-prey interactions is not factored into the Lima and Dill (1990) framework, nor it is factored into new conceptual frameworks for disgust [15,24], a form of parasite avoidance. Surviving while being fed on by parasites or micropredators opens up a broad range of responses that slow or stop feeding, or otherwise minimize its impact. Immune responses are a clear example of host responses made during parasite feeding that do not occur in predator-prey systems. Immune responses and other trait responses during parasite and micropredator feeding also cause non-lethal effects to individuals and broader ecosystems [5,21], making their inclusion in systematic examinations of non-lethal effects essential.

#### A general trait response framework

Although trait responses take diverse forms, all responses may be included in our proposed framework (Fig. 1). The framework is informed by a general consumer-resource model [27]. The general consumer-resource model features a single mathematical logic for characterizing the dynamics of all types of host-parasite and predator-prey systems (and other consumer-resource interactions, e.g. decomposer-carcass; plant-pollinator) by breaking down interactions temporally, during which individuals transition through discrete sequential states (Fig. S1). Prey and hosts transition through four states – susceptible, exposed, ingested, resistant (i.e. invulnerable; Fig. S1). For clarity, here we use the term ‘consumed’ in place of ‘ingested’. At the same time, predators and parasites transition through three states - questing, attacking, consuming (Fig. S1). Individuals transition between states following sequential biological processes: contact, attack failure and success, and feeding (further details of the model are provided in the supplementary material).

**Fig. 1.**
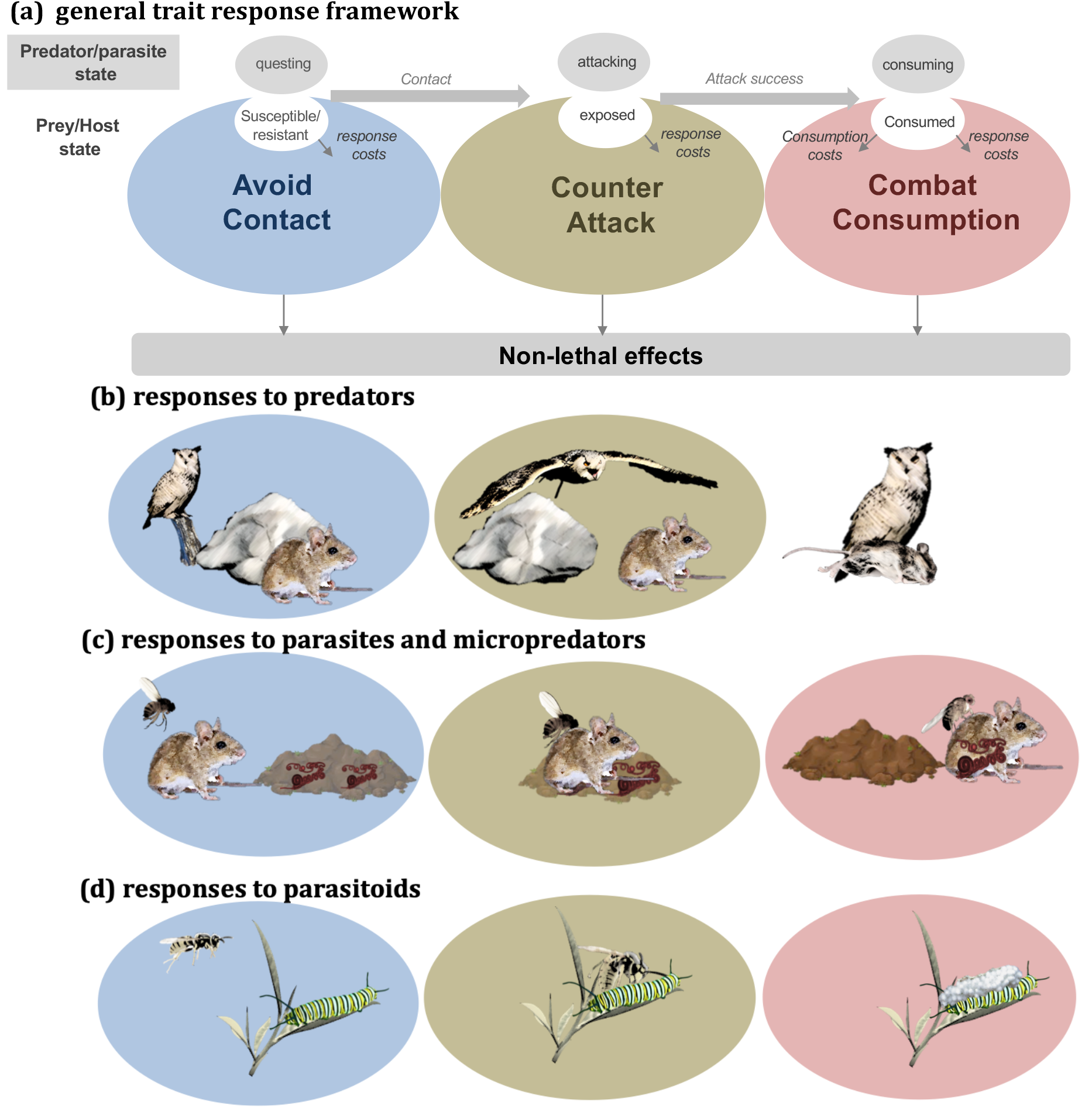
A general trait response framework and predictions. **(a)** Resources can mount three sequential responses to consumers, each with distinct effects on the interaction: avoid contact, counter attack, combat consumption. Examples of each type of response are listed in italicized text. Responses may be constrained by physical and sensory limitations, as well as trade-offs against other fitness-related activities (e.g. feeding and reproducing). The general framework can be tailored to specific types of consumer-resource interactions, such as interactions between field mice and **(b)** owl predators, **(c)** biting flies and infective nematodes (red worms in faeces), or **(d)** between caterpillars and parasitoid wasps. The lack of a combat stage in **(b)** illustrates that resources rarely respond during consumption by predators.

We distinguish three trait response classes in our framework that coincide with the biological processes driving predator-prey and host-parasite interactions: **avoid contact, counter attack,** and **combat consumption** (Fig. 1). All three responses can reduce the fitness costs of being consumed, albeit through distinct mechanisms and to varying degrees depending on their efficacy (i.e. the degree of change they cause to the interaction sequence). *Susceptible* prey and hosts may **avoid contact** with *questing* consumers before an attack. Effective avoidance reduces the rate that *questing* predators and parasites transition to *attacking,* and prey and hosts benefit from transitioning more slowly from *susceptible* to *exposed* states (Table S3). Prey and hosts that become *exposed* may **counter attack** to increase the probability that attacks fail. Countering-attack includes “fight or flight” responses, like hares sprinting to burrows when being chased by lynx, or tadpoles rapidly and erratically swimming when being attacked by trematode cercariae [12]. Finally, prey and hosts that are *consumed* (i.e. eaten or infected) and remain alive may **combat consumption**. Combating consumption shortens or slows predator and parasite feeding rates [i.e. ‘resistance’ in parasitology [19]], or lessens damage from the feeding [i.e. ‘tolerance’ in parasitology [20]]. Responses that reduce feeding include behaviours like social grooming by primates [28] and adaptive immune responses to parasitism [22]. Increasing tissue repair and protecting high-risk areas of the body from parasite feeding, as tadpoles do in response to trematodes [12], exemplify responses that combat consumption by reducing damage, without affecting feeding rates. Collectively, the combination of three trait responses with two consumer types results in six general pathways for non-lethal effects to arise.

#### Factors shaping the strength, duration, and frequency of trait responses

With the full range of trait responses classified and integrated into consumer-resource dynamics, we can now consider the conditions that determine which responses predators and parasites are likely to elicit, as well as how strong and frequent they are likely to be. The extent to which prey and hosts avoid, counter, and combat consumers depends first and foremost on their basic physical and sensory abilities [29]. For instance, tadpoles cannot physically leave ponds when predators are present. They may, however, reduce activity to avoid contact [30]. Individuals must also be able to detect consumer threats to respond to them. Prey and hosts use both visual and non-visual cues to detect predation and parasitism risk, making sensory limitations (e.g. sight, hearing, and smell) potential constraints on trait responses [31,32]. Impediments to either risk detection or the ability to act on detected risk should reduce the frequency of trait responses, or even preclude them.

Even when prey and hosts have the physical and sensory capacity to mount responses, trade-offs may mediate the frequency and strength of trait responses [29]. The ecology of fear provides ample evidence for the role of trade-offs in shaping trait responses to predators [33,34]. Whether via reductions in foraging, reproduction, or energy levels, fitness-related costs of mounting responses compete with the benefits of responses (i.e. the costs of not responding), making the frequency and strength that individuals exhibit trait responses a matter of economics [29]. An important point not recognized by the ecology of fear (or disgust) is that trade-offs may change over different phases of any given interaction. The frequency, duration, and strength of trait responses will therefore depend not just on their absolute costs but also on the relative costs compared to other possible responses. Exemplified by certain host-parasite interactions, avoiding contact may be more costly than becoming infected, potentially driving stronger and more frequent combat responses after becoming infected. The extent to which ‘fear’ and ‘disgust’ influence prey and host populations will depend on the feasibility to counter attacks and combat consumption. Phase-specific trade-off situations are accounted for in our framework through functions that link responses to state-specific mortality and fecundity rates, which expresses responses costs. Those costs are balanced by benefits of responses, expressed through response impacts on state transition rates, as described in the above section.

#### Predicting trait responses and non-lethal effects across different predators and parasites

The timing, frequency, and strength of trait responses in prey and hosts will also depend on traits of the predators and parasites, which leads to predictions for how trait responses and non-lethal effects may vary across the different types of consumers. Predators and parasites comprise distinct ‘consumer strategies’; that is, basic ways by which individuals feed on and impact other organisms [27,35]. One key difference involves the fitness consequences of predation and parasitism on individual prey or hosts. Predators and most parasitoids eliminate future reproductive success of their prey or hosts by killing them, and parasitic castrators do so by blocking host reproduction. In contrast, feeding by most other types of parasites and micropredators is not so deadly and does not completely eliminate future fitness gains. Given their great risk of consumption, we predict that predators, parasitoids, and parasitic castrators will elicit strong trait responses in their prey and hosts, resulting in overall strong non-lethal effects. Similar patterns may also emerge with other parasites with strong negative fitness impacts, such as highly virulent pathogens, or the micropredators that vector them. However, non-vectoring micropredators and less damaging parasites should elicit relatively weaker responses and weaker associated non-lethal effects. Differences in the fitness consequences of being consumed should then drive variation in non-lethal effects not just between predators and parasites, but also between different types of parasites, with some parasites being more similar to predators than to other types of parasites.

We can go further by considering the timing of prey and host death during interactions. Predators usually kill prey before or shortly after commencing to feed. This is not the case for most parasites and their hosts. Even parasitoids and castrators can have a substantial amount of feeding time before hosts are killed or castrated. Recognizing this basic difference in timing of prey and host mortality, or “reproductive death”, leads to two clear predictions for how non-lethal effects of predators and parasites differ. First, prey responses to predators will be constrained to avoiding contact and countering attack (Fig. 1b). Second, parasites and micropredators will elicit all three trait responses (Fig. 1c-d), resulting in more diverse non-lethal effects on hosts than prey of predators.

Integrating the above hypotheses leads to the prediction that, all else being equal, parasites and micropredators that entail high fitness costs should cause the strongest overall non-lethal effects from trait responses. This includes parasitic castrators, parasitoids, highly damaging pathogens, and the micropredators that vector them. The high fitness costs of feeding by these consumers should drive strong avoidance and counter-responses, resulting in strong non-lethal effects from those responses. Yet, by feeding on living organisms, these consumers also permit combat responses as a source of non-lethal effects, as evidenced in many hosts of parasitoids, parasitic castrators, and other parasites [5,36]. Collectively, we expect non-lethal effects from this triad of trait responses to supersede those arising solely from ‘fear’ responses (i.e. avoidance of contact and counter of attack) to predators. Exceptions may exist, such as some parasitoids that paralyze hosts during the initial attack, which can make hosts incapable of combatting consumption [36]. On average, however, our framework indicates that non-lethal effects will be strongest not solely from consumers that impose the highest risk of death, but rather from those that impose such strong costs while keeping the consumed host or prey alive.

### Studies to measure trait responses to predation and parasitism

We reviewed the literature that compares trait responses to predation and parasitism to assess support for our predictions. Specially, we compiled studies that directly compared trait responses to both predation and parasitism risk in a single resource species (see supporting material for details of literature search). The vast majority of studies we found measured trait responses in larval frogs (i.e. ‘tadpoles’, N = 106 entries across 13 studies, Table S1). Behavioural traits were most common, with activity level being the most reported trait (Table S1, Table S2). We did not find studies that measured immunological trait responses, likely because this is distinct to host-parasite systems. The studies spanned the following consumer strategies: solitary predators, trophically transmitted parasites, typical parasites, and pathogens (Table S1); in no cases were responses to parasitoids, parasitic castrators, micropredators, or social predators considered. Predator-induced trait responses were only measured during *questing* predator states, representing avoidance of contact, whereas measurements of parasite-induced responses included all three states: *questing* (N=9), *attacking* (N=5), and *consuming* (N=24). The limited data we found constrained our ability to make comparisons, although some general patterns did emerge.

Analysis of the data on tadpoles, the main animal group used as prey and hosts revealed considerable variation in the magnitude and direction of tadpole responses to predators (Fig. 2a), parasites (Fig. 2b), and their combination (Fig. 2c). However, on average and across all consumer and resource states, predator-induced trait responses were stronger than parasite-induced trait responses (Fig. 3a, Table S4). These patterns were evident after controlling for consumer state (comparing questing predators to questing parasites) (Fig. 3b), though they were weaker (Table S4), likely owing to lower power of the data subset. Distinguishing between trait response types (avoid, counter, combat) revealed that tadpoles did respond to parasites, but only by combatting parasites post-infection and at lower magnitudes than their responses to avoid predator contact (Fig. 3b, Table S4). Tadpoles also responded to the simultaneous presence of predators and parasites on average, and at similar magnitudes to their responses to predators alone (Fig. 3a). The strong tadpole responses to predators primarily represented reductions in activity levels (Fig. S7a, Table S4), and they were most evident when measuring group-level responses as opposed to individual-level responses (Fig. S7b, Table S4). Across the host-parasite interactions studied, responses did not differ between pathogens and trophically-transmitted parasites, the two parasite strategies investigated (Table S3).

**Fig. 2.**
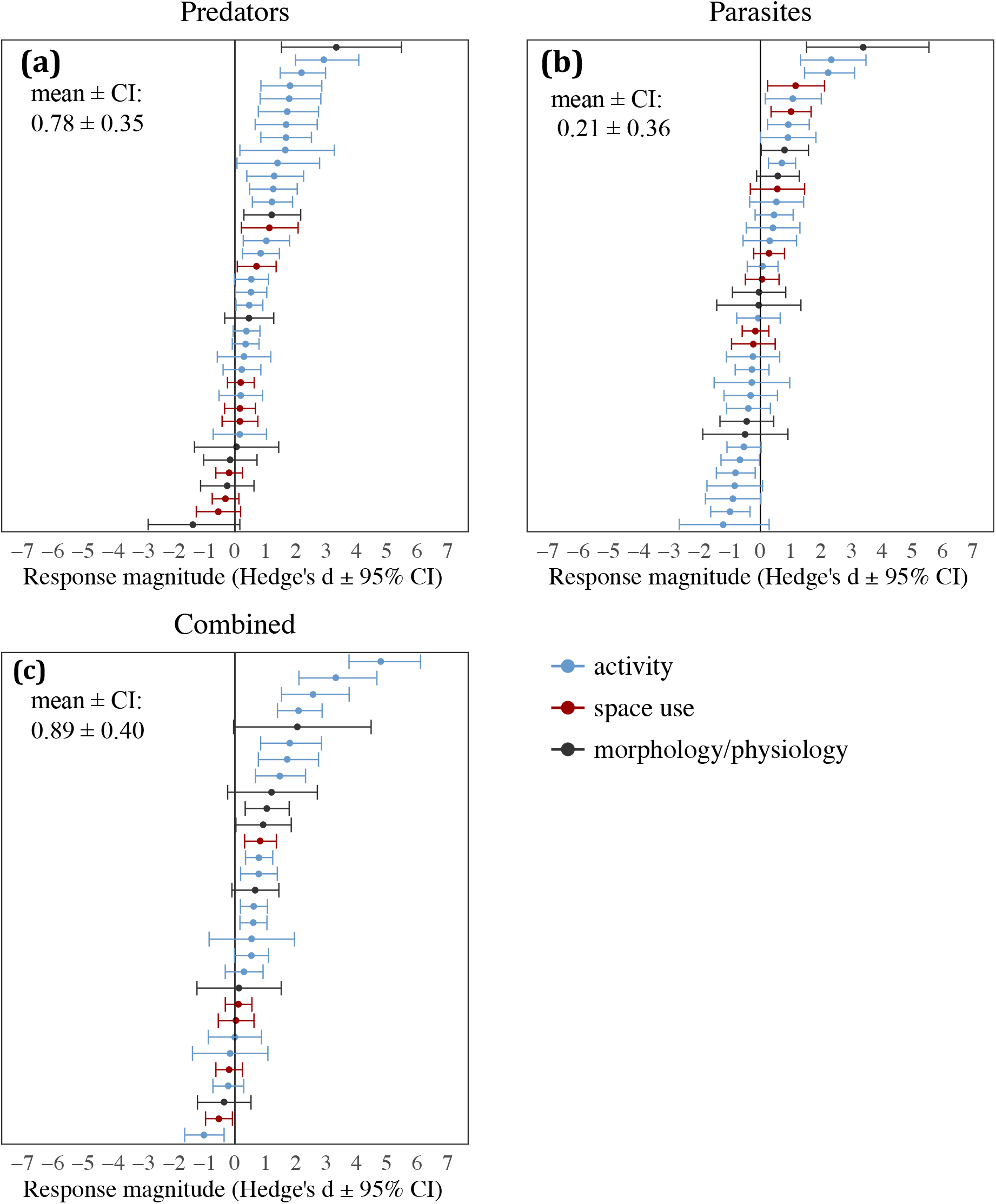
Forest plots of effect sizes used in the meta-analysis. The distribution of effect sizes for responses elicited by the presence of **(a)** predators, **(b)** parasites, and **(c)** their combined presence resulting from resource adjustments in activity (blue), space use (red), and morphological/physiological traits (grey). Error bars denote the 95% confidence intervals.

**Fig. 3.**
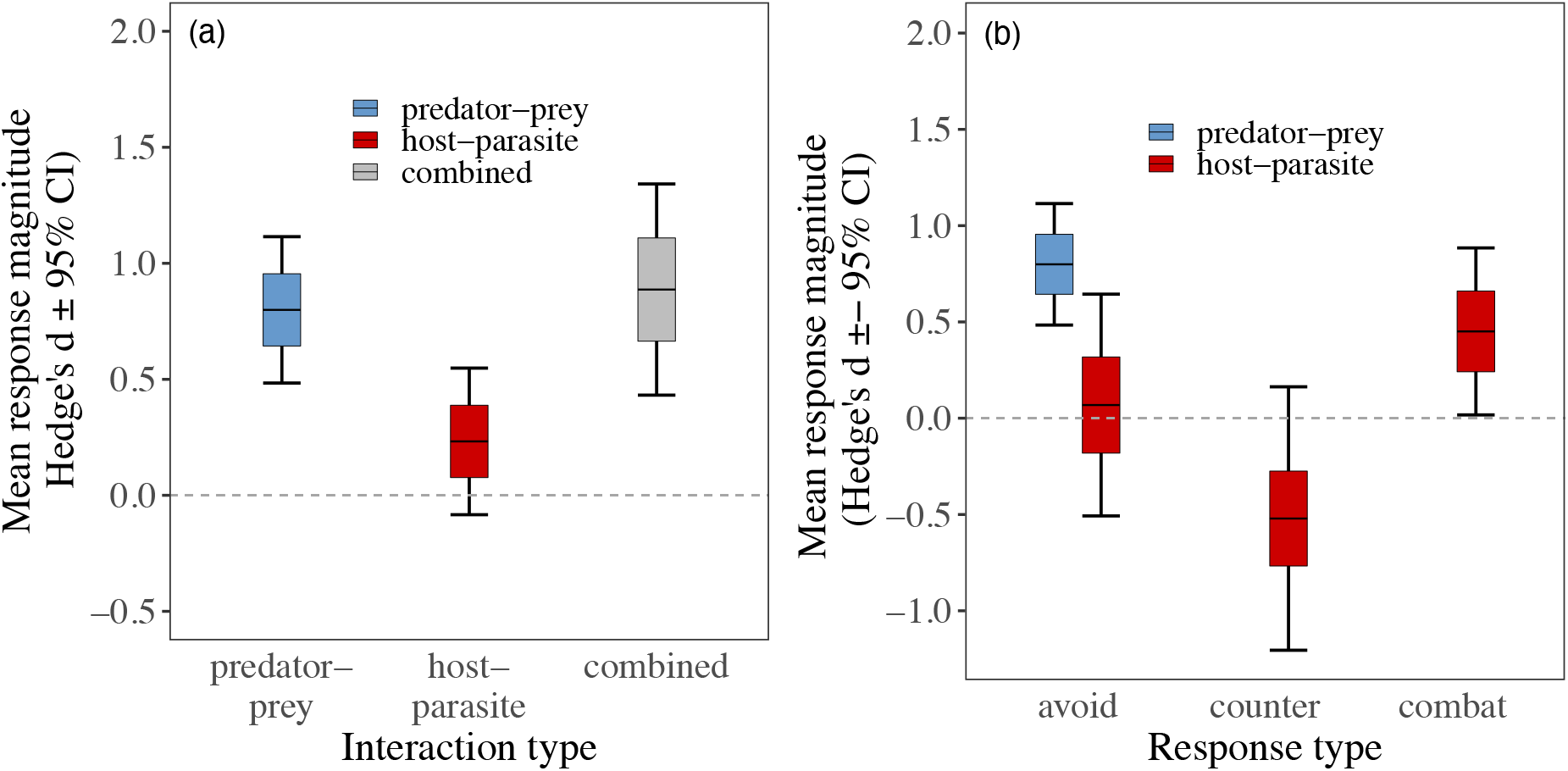
Relative magnitude of responses to predation vs. parasitism. **(a)** The estimated mean magnitude of trait responses to predation cues (blue), parasitism cues (red), and both cues (grey). **(b)** Mean trait responses to predators (blue) and parasites (red) when distinguishing by the type of trait responses, as defined in our framework. Only avoidance responses to questing predators were found in our literature review, likely owing the low probability of surviving attack or consumption by predators. Responses to the combined presence of predators and parasites are not shown in **(b)** because only one study with this treatment had predators and parasites in the same state. Lines denote the mean response magnitudes, boxes denote the standard error of the mean, and error bars denote the 95% confidence intervals.

## Discussion

Our framework reveals that non-lethal effects from predation and parasitism are a function of diverse and potentially interacting responses by prey and hosts, exhibited at different time points in consumer-resource interactions. The current literature on non-lethal effects provides a very limited portrayal of trait responses, which restrained our ability to test the predictions that emerged from the general framework. Current comparative data comprise brief snapshots of interactions that do not fully capture the temporal dynamics of trait responses (and their non-lethal effects). Longitudinal data that tracks individuals through all interaction phases could provide insight into how non-lethal effects accrue from multiple trait responses. Longitudinal data could also detect interactive effects between trait responses. For example, hosts that invest heavily in immune defences may exhibit weaker avoidance of contact with parasites, particularly if avoidance conflicts with feeding, reproducing, and other fitness-related activities. The growing literature on ‘fear’ and ‘disgust’ of parasites, which focuses exclusively on certain forms of parasite avoidance, will benefit by being contextualized within the broader suite of trait responses available to hosts.

Trait response data on tadpoles, although limited, underscore the importance of distinguishing nuances in the timing of trait responses. Pooling all trait response measurements suggested that parasites generally did not elicit changes in tadpole activity levels as did predators. However, accounting for response timing revealed significant changes in activity levels that were confined to post-infection periods of interaction. Adult amphibians show similar tendencies to combat infections rather than avoid them [37]. Combat responses appear to be a primary trait response to parasitism in this group of animals, whose role in non-lethal effects can most fully be estimated by treating the responses as separate from those made earlier in interactions.

The post-infection combat responses apparent in tadpoles are particularly notable considering that immune responses were not factored in. Immune responses are a very common form of combatting parasitism that can be exhibited for prolonged periods. The combined non-lethal effects of tadpole parasites were therefore likely much stronger than the analysed data suggest. Additionally, non-lethal effects of parasitism could have occurred from host phenotypic changes caused by parasite manipulation [25,38], and even directly from parasite feeding independent of defensive responses. For instance, general energy drain or direct tissue damage caused by parasite infection can substantially impact host performance [39,40]. The diverse sources of non-lethal effects of parasites not relevant to prey collectively could have rivalled in magnitude the stronger predator avoidance observed in tadpoles.

Accounting for distinct predator and parasite strategies led to trait response predictions that did not align with the predator versus parasite dichotomy. Pinpointing the sources of variation in non-lethal effects therefore demands a functional approach rather than a taxonomic approach to distinguishing predators and parasites [35]. Parasitoids, and parasitic castrators in particular, could offer rewarding insights because they share functional characteristics with both predators and parasites. Avoidance, counter, and combat responses to parasitoids are well-documented [36], but how their frequency and strength compare with responses to predators and less debilitating parasites is unknown. The latter can also be said for parasitic castrators. Given the high fitness consequences of infection from parasitoids and castrators, together with infections that are not immediately lethal, their combined non-lethal effects may very well be the strongest of all predator and parasite functional groups.

Regardless of how individuals respond to predators and parasites alone, risks of predation and parasitism in real ecosystems rarely occur in isolation. Future research could apply our framework to investigate the additive and interactive non-lethal effects of simultaneous exposure to predators and parasites. Our analysis of the tadpole data suggests that responses to simultaneous exposure are non-additive, perhaps owing to the prioritization of responses to the more severe threat (Fig. S5). Although not a focus of this review, evidence for increasing predation of parasitized prey [41–43], and increasing parasitism in predator-rich environments [44], provide further indication that predators and parasites interact to impose non-lethal effects in non-additive ways. Fewer studies have considered single responses that defend against both predators and parasites. Nevertheless, there were several cases where tadpole responses to predators and parasites responses were in the same direction (i.e. a reduction in the trait expression). Trait responses that effectively deter both predators and parasites may mitigate the non-lethal effects incurred from the essential task of defending oneself against being eaten.

## Conclusion

Whether through fear or through infection, consumers elicit costly trait responses in their resources that give rise to non-lethal effects at the level of individuals, with consequences for communities and ecosystems more generally. A general consumer-resource model helped us to develop a framework for systematically comparing trait responses to various types of predators and parasites. Different types of predators and parasites should elicit different trait responses, and therefore, have different non-lethal effects, given differences in consumer strategies that influence when and how strongly they impact prey and hosts. However, many predator and parasite strategies have not yet been tested comparatively. Expanding research of non-lethal effects to regularly consider different predator and parasite strategies sets the foundation for exploring how non-lethal effects manifest in the multi-dimensional food webs found in real ecosystems.

## Acknowledgements

We thank Gordon Research Conferences and Andy Sih, chair of the 2016 meeting “Predator-Prey Interactions”, for providing the forum that inspired this work; the EGLIDE group (Amy Pedersen, Amy Sweeny, Saudamini Venkatesan, Dishon Muloi, Alexandra Morris, Shaun Keegan and Kayleigh Gallagher), the EEGID group at University of Liverpool (Mike Begon, Greg Hurst, David Montagnes, Mark Viney, Steve Parratt), the Garner lab at the Zoological Society of London (Trent Garner, Chris Owen, Lola Brooks, Stephen Price, Goncalo Rosa, Bryony Allan), and Chris Carbone for their comments on earlier versions of the review; and Daniel Noble, Raj Whitlock, and Wolfgang Viechtbauer for advice on the meta-analysis. Any use of trade, product, or firm names in this publication is for descriptive purposes only and does not imply endorsement by the U.S. Government. This manuscript benefited from NSF Ecology of Infectious Diseases grant (OCE-1115965) and a grant from the Natural Environment Research Council UK (NE/N009800/1).

## Author Contributions

This review was broadly conceived during a 2016 Gordon Conference group meeting on predator-prey interactions. DRD and EM organized initial planning and idea development. DRD performed the systematic review and meta-analysis with input from the other authors. DRD, KL, RH, and AF constructed the framework and drafted the manuscript. All authors contributed to the revision process and ideas expressed in the final piece.

## Data accessibility statement

The data and code supporting the review and data analyis are archived in Dryad and will be made publicly available should the paper be accepted after peer review.

