## supplementary material for "Broadening the ecology of fear: non-lethal effects arise from diverse responses to predation and parasitism"

### Detailed methods for the systematic review of trait response literature

The goal of the systematic review was to (a) assess the breath of empirical studies to directly compare trait responses to predators and parasites, and (b) seek support for the predictions emerging from our general trait response framework.

*Data collection and inclusion criteria*

We searched for literature using Google Scholar and Web of Science, using combinations of terms describing consumers (consumer, predat*, parasit*, anti-parasit*, anti-predat*, infect*, natural enemy), resources (host, prey, resource), traits (behavio*, activity, space use, refuge use, feeding, development), and effects (non-lethal, sub-lethal, risk, trait-mediated, trait response, indirect). We initiated our search on 10 January 2017 and continued to monitor the literature until the time of manuscript submission. We considered journal articles that met three criteria:

1. measured traits associated with defense in an animal exposed to a predator cue. Cues included media (e.g. water) used by a predator (Han et al. 2011; Szuroczki & Richardson 2012; Marino et al. 2014), a consumed conspecific (Rohr et al. 2009), a predator kept behind a barrier (e.g. mesh cage) (Thiemann & Wassersug 2000; Parris & Beaudoin 2004; Raffel et al. 2010; Preston et al. 2014), or an unrestrained predator.

2. measured the same trait response (in the same resource species and life stage, but not necessarily the same individuals) to 1) a parasite cue, such as media previously containing infected hosts or infective stages (Preston et al. 2014) or direct exposure to parasite infective stages, or 2) in animals infected with parasites.

3. had a control, wherein defense traits were measured in individuals that were not exposed to either a parasite or predator cue.

We did not include studies of trade-offs between predation and parasitism that used infection (in the case of predator effects) or predation (in in the case of parasite effects) as response variables because different traits were measured between "predator" and "parasite" treatments, and therefore were not comparable.

**Analysis of the tadpole data**

Studies included in our systematic review were limited but nevertheless provided adequate data for performing an initial quantitative analysis of how the magnitude of trait responses differ between predators and parasites. We used standard meta-analytic approaches to estimate mean magnitudes and heterogeneity of responses, as well as the factors influencing them. We point out however that we did not intend for this to be an exhaustive meta-analysis of all available trait response data. Given that the vast majority of included studies used larval and amphibians, we focused our analysis on this group of animals. Repeating the analyses with the full dataset did not qualitatively change our findings.

*Data extraction*

We used raw data from studies when available. Otherwise, we used the *digitize* package in R to extract data from figures (Poisot 2011; R Core Team 2013). We recorded the number of replicates as well as the mean and variance (SE or SD) of the response variables separately for controls, predator treatments, and parasite treatments. Studies often included a treatment containing both a predator and parasite cue (hereafter referred to as 'combined'). We recorded these treatments as separate entries, with independent sample sizes, means, and variances. For studies that measured responses in multiple resource species and/or for multiple traits, we treated each species and trait as a different entry. When data were recorded across multiple days, we pooled data to calculate a single response mean and variance. We also pooled data from any additional treatments that varied factors such as consumer density (Raffel *et al.* 2010; Preston *et al.* 2014) or exposure frequency (Preston *et al.* 2014), and again calculated the response mean and variance for predator, parasite and, when applicable, combined treatments.

*Factors influencing responses*

For each entry we distinguished the interaction type ('predator-prey', 'host-parasite', or 'combined') and recorded the following variables: a.) trait type – the focal trait measured to gauge responses, classified here as 'activity' (e.g. proportion of time moving, grooming, etc.), 'space use' (e.g. proportion of time in a refuge, distance to consumer cue, etc.), or 'morphology and physiology (e.g. mass, development rate), b.) analysis scale –whether studies used a trait value of individuals ("individual level" e.g. proportion of time spent in refuge) or proportions of individuals in group that exhibited the focal trait ('group-level'. e.g. number of individuals that used a refuge) as the unit of measure; c.) consumer presence – whether the consumer was present in treatments ('direct') or treatments used cues such as media used by consumers or ingested conspecifics to elicit responses ('indirect'), d.) the genus and species of the consumer and resource, e.) the state of the consumer when the resource trait was measured (questing, attacking, consuming), and f.) the trophic strategy of the consumer (Lafferty & Kuris 2002). See table S1 for a complete breakdown of how the type and scale of trait responses were classified.

For predator treatments in which non-visual cues were used or in which predators were kept behind a barrier, we considered predators to be questing and resources to be in susceptible states. Determining the states of parasites during host responses was less straightforward. We used the following criteria to determine parasite states at the time when response traits were measured in hosts:

questing parasites/susceptible hosts: Infective parasite stages were kept separate from resources (e.g. via a screen with mesh fine enough to prevent parasite escape);

b.) attacking parasites/exposed hosts: traits were measured immediately after exposing hosts to unrestrained infective stages;

c.) consuming parasites/ingested hosts: traits were measured 24 hr post-exposure (infective stages of many commonly studied parasites, such as trematode cercariae and fungal spores rarely live longer than 24 hrs), or if infections were confirmed in hosts prior to taking the measurements.

Experiments evaluating effects of trophically-transmitted parasites at times had two controls: absence of infective parasite stages but the presence of the uninfected alternative hosts and absence of both infective parasite stages and alternative hosts. For example, to assess responses to amphibian trematodes with snail alternative hosts, caged uninfected snails as well as the absence of snails were used as controls for treatment groups with caged infected snails (Rohr *et al.* 2009; Preston *et al.* 2014). For these studies, we considered the uninfected alternative host treatment as the control group to account for potential influences of alternative host presence. If studies exposed hosts to free-living infective parasite stages (Thiemann & Wassersug 2000; Han *et al.* 2011; Preston *et al.* 2014), data from the media-only treatment were used for the control.

*Effect Sizes*

We used the standardized mean difference that accounted for unequal variances and small sample sizes (i.e. *Hedge's d*) as the measure of effect sizes (Koricheva *et al.* 2013). Here, *Hedge's d* represents the mean difference in trait values of resources in treatment groups versus those of resources in control groups, expressed in units of standard deviations. *Hedge's d* factors in sample sizes, mean trait values, and variances in treatment and control groups, respectively. We used the number of replicates as the sample size and the pooled standard deviation as the unit of variance. Most studies reported variances as standard errors of means, in which case we calculated the standard deviation as: SE * SQRT(N), where SE is the standard error and N is the sample size (number of replicates in our case).

We manually computed effect sizes and compared our values against those obtained from the *escalc* function of the metafor package (version 1.9-9) in R (Version 3.3.2) (R Core Team 2013) as an accuracy check. We calculated the effect size so that positive effects denoted reductions in trait values (e.g. reduced activity level), except for measurements of time in a refuge or distance from a consumer cue. For consistency, we reversed the sign of measures of space use so that positive effect sizes would denote increases in the use of low risk spaces (e.g. refuges; Table S2). Any influence of this sign reversal was accounted for by including trait type in our models (see below).

*Statistical Analyses*

We estimated mean response magnitudes for each interaction type ('predator-prey', 'parasite-host', 'combined') and other factors of interest (trait type, analysis scale, consumer presence) by running a series of multi-level generalized linear mixed models (GLMMs) using the rma.mv function from the *metafor* package (Viechtbauer 2010) in R (studio version 1.0.136) (R Core Team 2013). Since data on responses to predators were available for questing states only (see results), we also compared response magnitudes to questing predators and parasites only. We used Hedge's *d* as the response in all models, and we used a Gaussian error structure because responses were normally distributed in all cases. All models also included entry as a random effect, nested within a third-level random effect of study because the contribution of multiple entries by single studies could create non-independence of the data (Konstantopoulos 2011). We did not control for consumer or resource phylogeny because of the low taxonomic diversity of our sample (Table S1). We used variance-covariance matrices as the sampling variances for models comparing responses across different interaction types to account for their multi-arm designs (designs that compared predator and parasite treatments to a single control; Gleser & Olkin 2009). For the analyses that only used parasite data we used the sampling variance, calculated with the *escalc* function in *metafor*, since these data comprised a single treatment group per control group. We also computed the $I^2$ statistic to assess how much of the total variance in response magnitudes were explained by between-study vs. between-entry heterogeneity (Higgins & Thompson 2002).

To test the effects of interaction type and other factors of interest on mean response magnitudes, we compared GLMMs including factors of interest as fixed effects with models omitting factors, using likelihood ratio tests against a chi-square distribution as a measure of significance. Confidence intervals (CI) were estimated from restricted maximum likelihood, and response magnitudes were considered significantly different from zero if the confidence intervals did not include zero. As a consistency check for the results of likelihood ratio tests, we also compared models and measured factor importance using information theoretic approaches.

*Publication Bias*

The specificity of the targeted studies used to compile our dataset, combined with their inclusion of multiple treatments and effect sizes (predation, parasitism, combined), should have minimized publication bias. Nevertheless, we assessed the likelihood of publication biases in the data by performing rank correlation tests (‘*ranktest*’ function in Metafor) and Egger’s regression tests (‘*regtest*’ function Metafor) for the three versions of the dataset used to construct our models: full dataset, exposure to questing consumer stages subset, parasite exposure subset. In addition, we estimated Rosenthal’s fail-safe N computation using *metafor*, which tests the robustness of effect size estimates using the number of studies with null results that are needed before Type I errors can be significant (Rosenthal 1979).

**Results**

*Summary statistics:* Tadpole responses to predators, parasites and their combined presence varied across studies (Fig. 2 in main text). Of the total variance, 85.5% came from heterogeneity in true effects between entries. Between-study heterogeneity was negligible (<0.01%), indicating that methodological differences between studies were not major contributors to observed variation response strengths. Predators elicited significant responses in 51% (20 of 38) of cases, all of which were positive (i.e. reduction in trait value; Fig. 2a). Parasites elicited responses in 26% of cases (10 of 38), 10 of which were positive and 3 of which were negative (Fig. 2b). Responses to the simultaneous presence of predators and parasites elicited significant responses in 53% of cases (14 of 30), and most were positive (positive effects = 14, negative effects = 2; Fig. 2c). All negative responses to parasites were elicited exclusively by attacking parasite stages and represented increases in activity levels.

*Factors influencing the average strength of trait responses:* Table S4 reports the outputs of models run to predict the factors driving trait responses. Briefly, likelihood ratio tests indicated that average response strengths were strongly influenced by interaction type, analysis scale, and their interaction (dropping their interaction reduced goodness of fit, Table S4). When pooling all data, predators and their combined presence with parasites elicited strong responses on average, but parasites alone did not. Strong responses were detected group-level measures (Fig. S7), such as the proportion of individuals in a group that were moving at a given time, rather than individual measures, such as the proportion of time that individuals spent moving. The type of trait measured had lesser of a general influence on response strengths, though most significant responses were reductions in activity levels, likely because that was the most common trait measured (Fig. S7). Controlling for consumer state by looking only at questing predators and parasite showed again that predators elicited responses on average, but parasites again did not. However, examining only parasite data and factoring parasite state into the models revealed that parasites did elicit responses in their hosts on average (dropping parasite state from models reduced goodness of fit, Table S4), but only by consuming states (i.e. post-infection).

*Multi-model inference:* Results corroborated the likelihood ratio tests. The best performing model according to AICC ranking included interaction type, analysis scale, and trait type as fixed effects (Table S6). Interaction type was the most important factor - it was included in all models within 2 AICc values of the top model, followed by trait type, which also had an importance ranking of over 80 percent (Fig. S3).

*Publication bias:* Funnel plots indicated a skew toward positive effect sizes for the three models used in our analyses (Fig. S4). Rank correlation tests and Egger's regression tests indicated potential publication bias for the models using the full dataset (rank correlation: tau = 0.237, $p < 0.001$; Egger's: z = 3.04, p = 0.002) and the subset of only responses to questing consumers (rank correlation: tau = 0.25, p = 0.01; Egger's: z = 2.69, p = 0.007). However, Rosenthal's fail-safe numbers were larger than the threshold level required for our analysis (cumulative: 9746, questing: 1940), indicating that the effect sizes of all models were robust to publication bias. No publication bias was detected for the models of responses to parasites only (rank correlation: tau = 0.10, p = 0.368; Egger's: z = 1.19, p = 0.254; fail-safe number: 80).

### Figures and Tables

**Effects of trait responses on the consumer**
-Increased mortality
-Return to questing state (predators only)
-Reduced attack opportunity and success

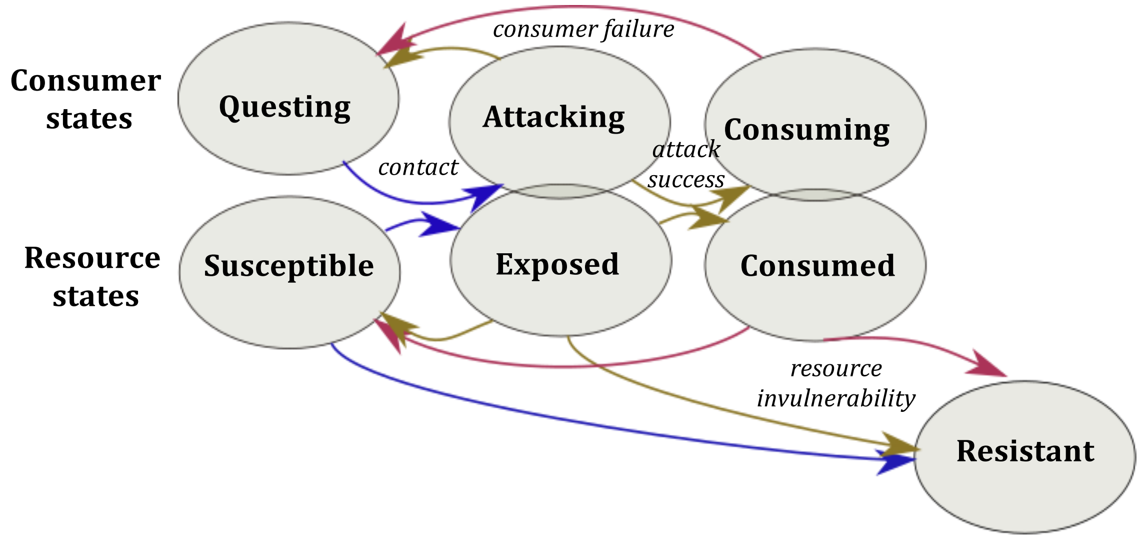

**Effects of trait responses on the resource**
-Reduced mortality
-Return to susceptible state
-Temporary or permanent invulnerability
-Reduced exposure and feeding rate

**Fig. S1. Effects of trait responses on consumers and resources.** Responses alter biological processes driving consumer-resource dynamics (contact, attack success, death, invulnerability), and in turn have distinct effects on the interaction. Blue arrows denote processes impacted by resources avoiding contact with consumers. Brown arrows denote processes impacted by resources countering consumer attacks. Red arrows denote processes impacted by resources combatting consumers during feeding.

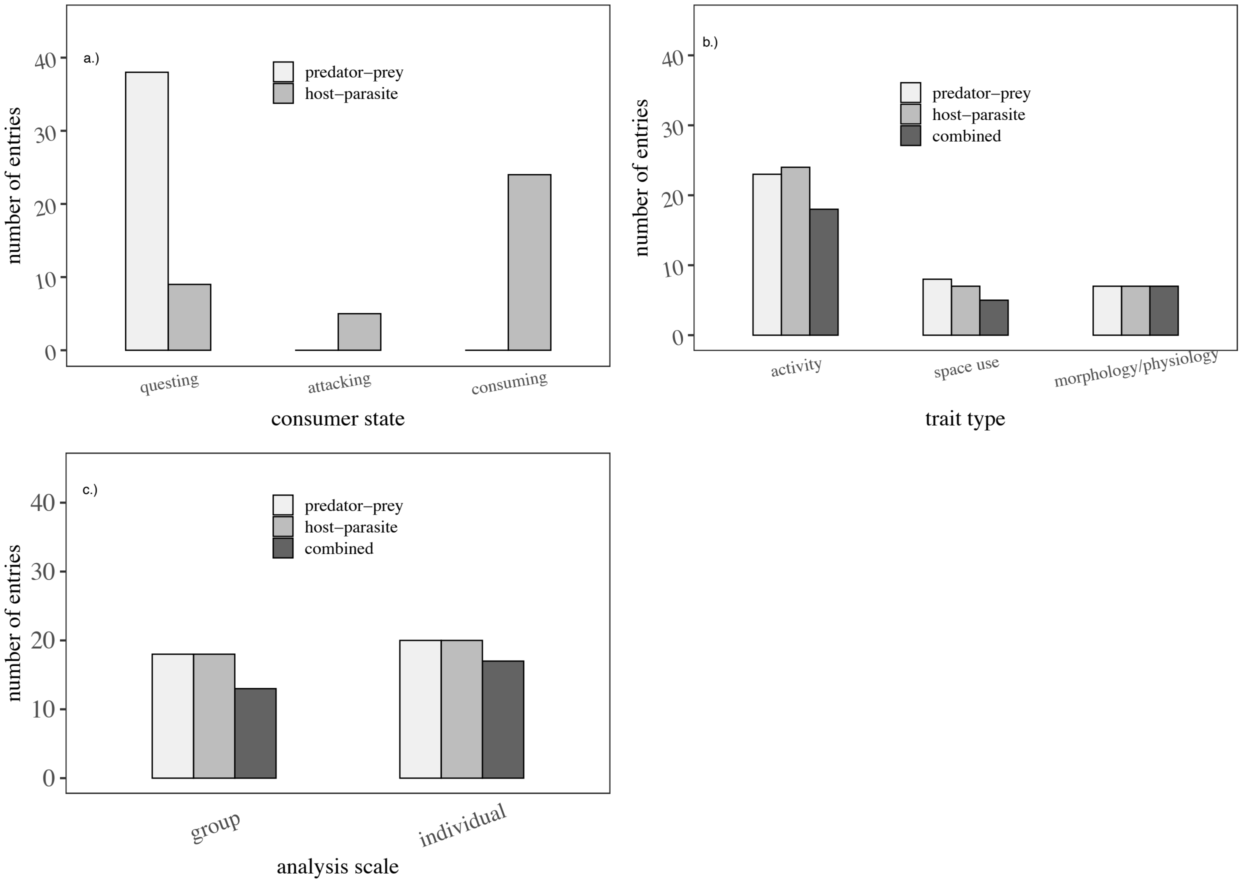

**Figure S2.** The number of entries used in the meta-analysis to calculate non-lethal effects resulting from prey or host responses to predation (blue) and parasitism (green). Parasite entries spanned all three consumer states, while predator entries focused on questing states (a). Entry numbers are also provided for the focal factors in our analysis, including (b) the type of trait measured, (c) and analysis scale – whether individual or group level responses were measured.

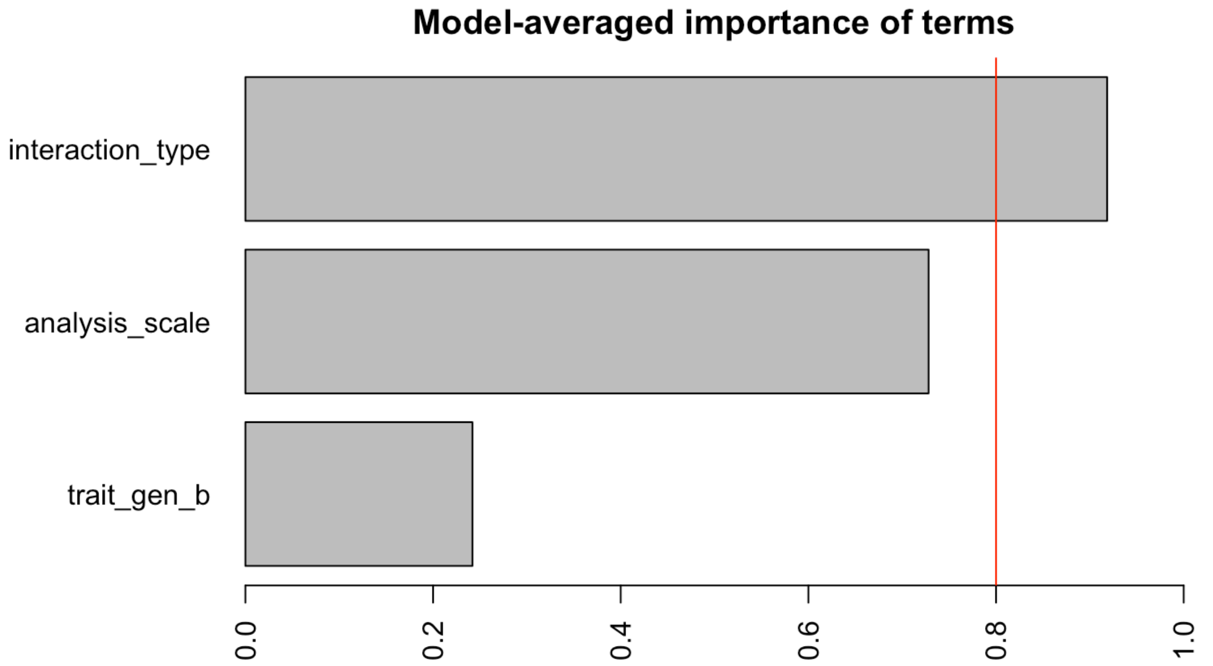

**Figure S3.** The model-averaged importance of the fixed effects included in models used for multi-model inference. The red line denotes an 80% importance, which is often used to distinguish important versus unimportant factors in explaining responses.

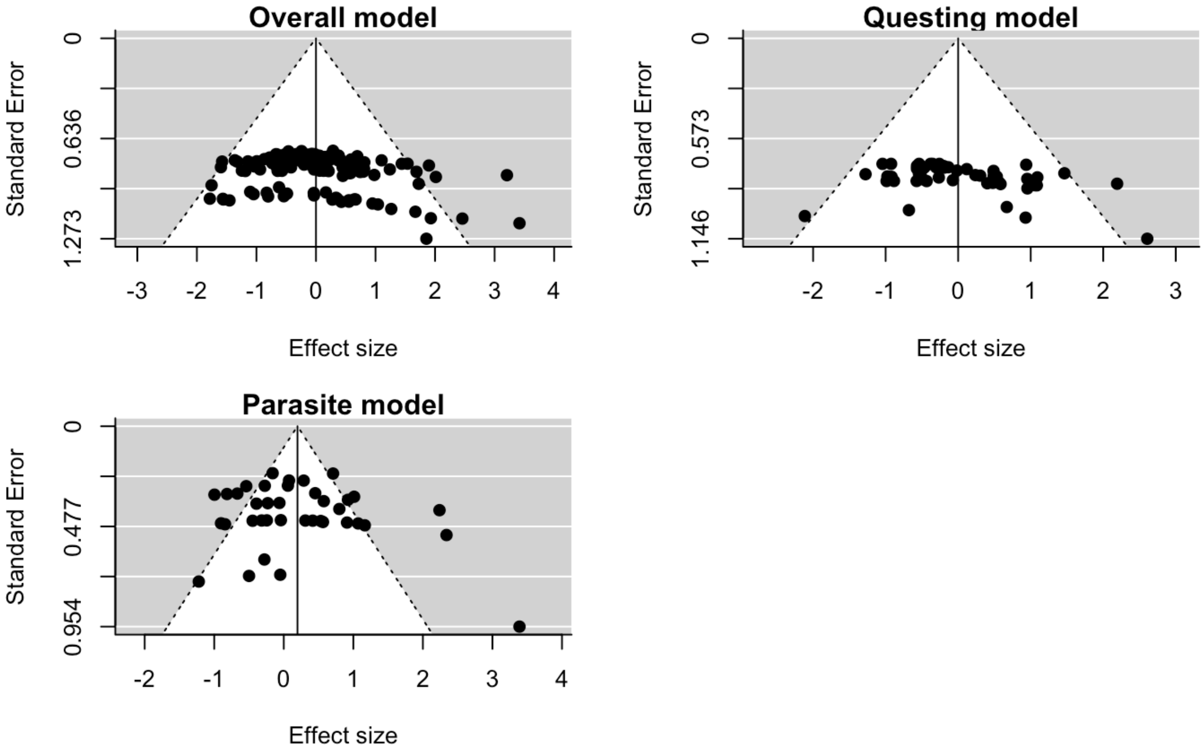

**Fig. S4**. Funnel plots for the three general model structures used in the meta-analysis. The cumulative model included the full dataset, while the questing and parasite models used a subset of the dataset pertaining specifically to consumers that were confined to questing states or were parasites, respectively. Asymmetries in point distributions about zero on the x-axis are indicative of publication bias.

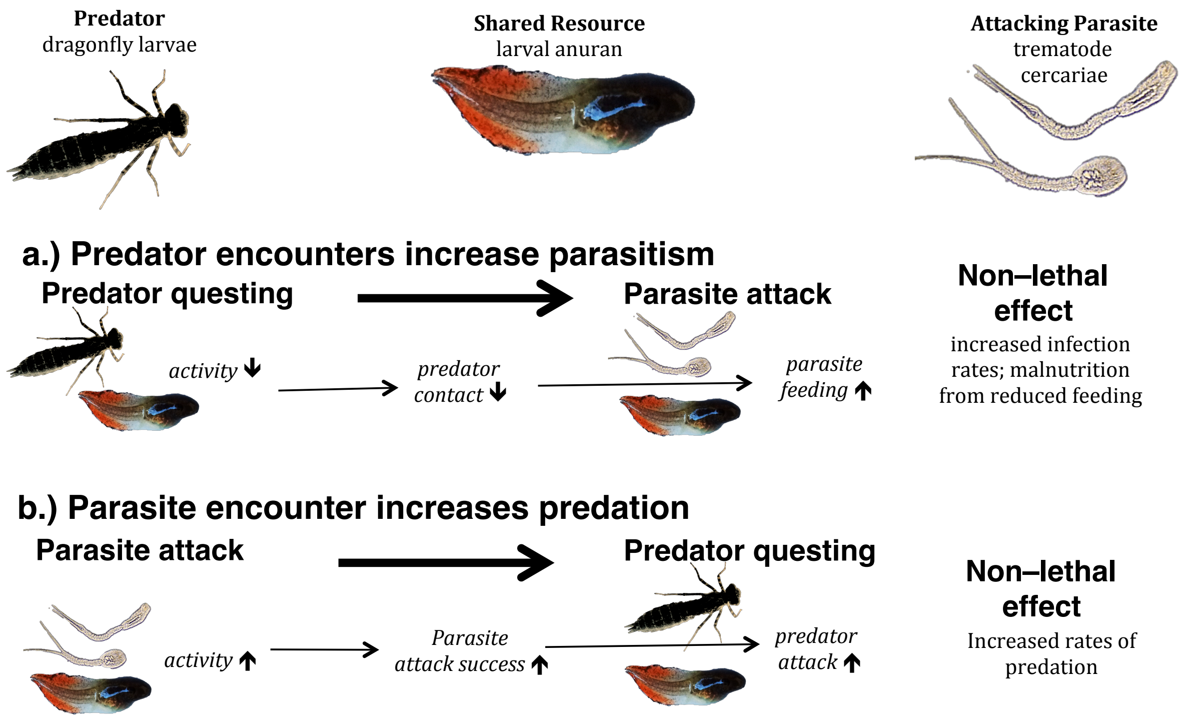

**Fig. S5. Scenarios where trait responses to predators and parasites interact across different phases of interactions.** Larval anurans are a shared resource of predatory dragonfly larvae (*Anax spp.*) and trematode parasites, and both consumers elicit behavioural changes in larval activity. Differences in the timing and direction of those changes suggests that priority effects may be important in determining interactive non-lethal effects of these consumers. For example, **(a)** initial exposure to questing dragonfly larvae strongly reduces tadpole activity, which successfully reduces contact with predators, but also likely reduces trematode attack failures, or **(b)** strong increases in tadpole activity in response to attacking trematodes results in greater trematode infection failure, but may also increase contact with questing dragonflies.

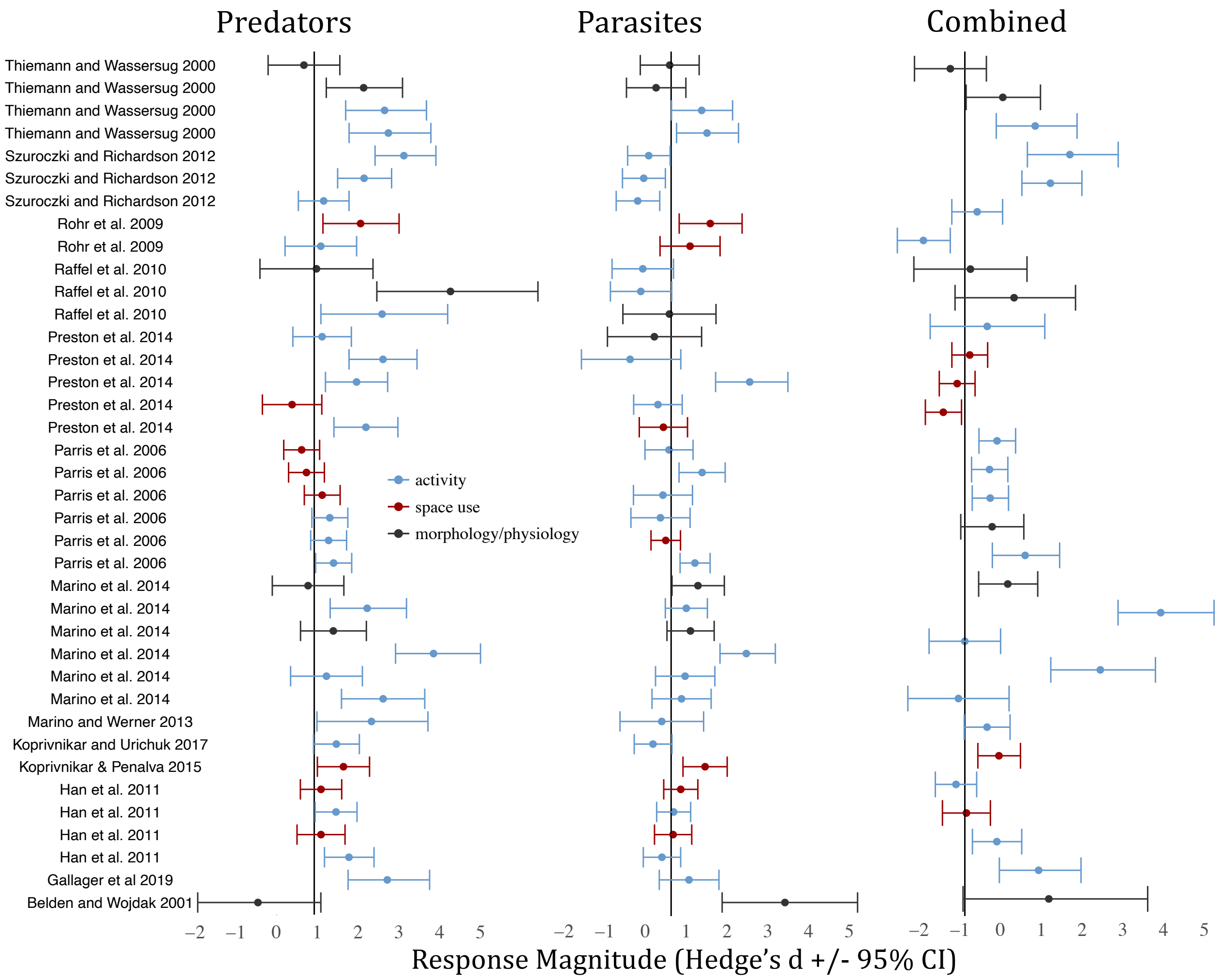

**Fig. S6. Forest plots effect sizes used in the meta-analysis, sorted by study.** The distribution of effect sizes for responses elicited by the presence of predators (left), parasites (middle), and their combined presence (right) resulting from resource changes in activity (blue), space use (red), and morphological/physiological traits (grey). Effect sizes are sorted by study to facilitate investigation of the data on a study by study basis. Studies may have contributed multiple entries by considering multiple traits, consumer species, or resource species.  Error bars denote the 95% confidence intervals.

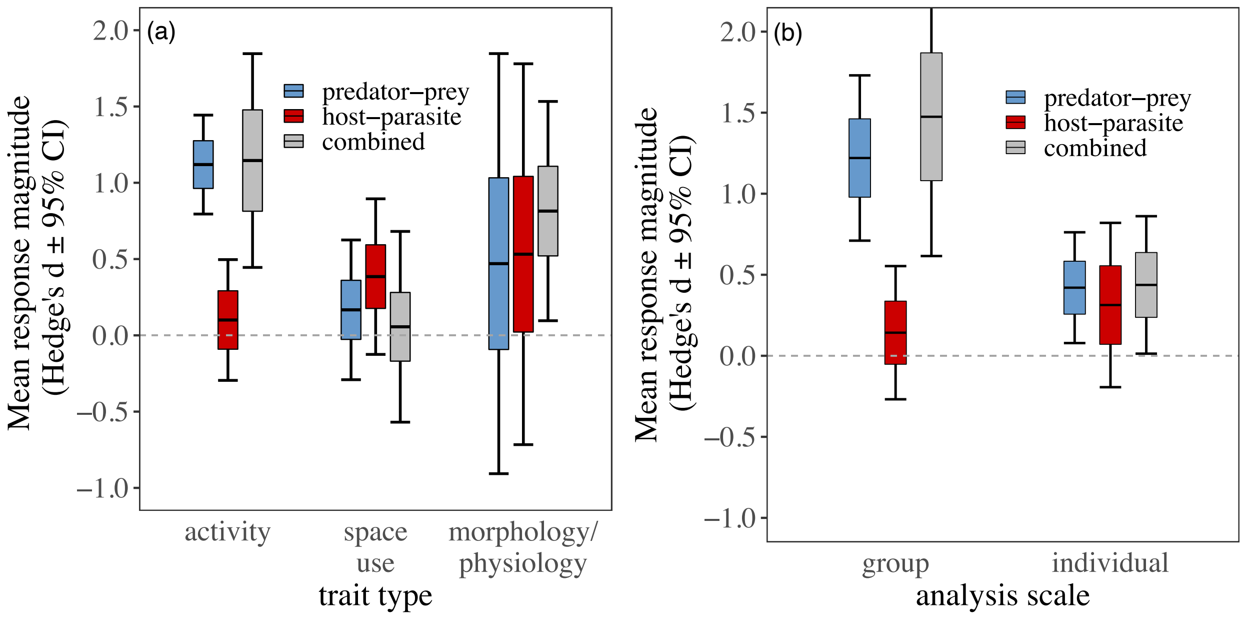

**Fig. S7. Magnitudes of responses to predation vs. parasitism depend on trait type and analysis scale.** Estimated mean trait response magnitudes to predation cues (blue), parasitism cues (red), and both cues (grey), broken down by **(a)** the type of trait measured and **(b)** the scale at which trait responses measured. Lines denote the mean response magnitudes, boxes denote the standard error of the mean, and error bars denote the 95% confidence intervals.

**Table S1: Direct empirical testing of non-lethal responses to predation and parasitism.** A list of studies that directly compared responses to the presence of predators and parasites by animals serving as their resource. Several studies were excluded from our meta-analysis owing to limitations in data reporting. (See attachment)

| Focal trait | trait type | analysis scale | Positive effect interpretation | effect size measure |
|---|---|---|---|---|
| time spent moving | activity | individual | less time spent moving | hedges *d* |
| number of lines crossed | activity | individual | fewer lines crossed | hedges *d* |
| time spent swimming | activity | individual | less time spent swimming | hedges *d* |
| time spent crawling | activity | individual | less time spent crawling | hedges *d* |
| time spent grooming | activity | individual | less time spent grooming | hedges *d* |
| time spent turning | activity | individual | fewer turns | hedges *d* |
| proportion of observations that resource was moving | activity | individual | fewer observations of movement | hedges *d* |
| proportion of time in foraging zone | space use | individual | less time foraging | hedges *d* |
| proportion of individuals moving | activity | group | fewer individuals moving | hedges *d* |
| number of individuals at water surface | space use | group | fewer individuals at surface | hedges *d* |
| number of times refuge was used | space use | individual | more time spent in refuge | hedges *d* * (-1) |
| distance to cue | space use | individual | maintained larger distance to cue | hedges *d* * (-1) |
| distance to consumer barrier | space use | individual | maintained larger distance to consumer barrier | hedges *d* * (-1) |
| vertical migration distance | space use | individual | higher distance climbed | hedges *d** (-1) |
| time in foraging habitat | space use | group | less time in foraging time | hedges *d* |
| proportion of individuals moving | activity | group | fewer individuals moving | hedges *d* |
| number of individuals at water surface | space use | group | fewer individuals at surface | hedges *d* |
| CORT level | morphology/physiology | individual | reduced stress levels | hedges *d* |
| development stage at end of experiment | morphology/physiology | individual | less advanced development stage | hedges *d* |
| mass | morphology/physiology | individual | reduced mass | hedges *d* |
| body length | morphology/physiology | individual | reduced length | hedges *d* |

**Table S2:** A list of traits measured by studies to estimate non-lethal effects of predators and parasites. Positive effects denoted reductions in trait values (e.g., reduced activity level or mass), except in measures of space use that measured time in a refuge or distance from a consumer cue (e.g. positive effect would mean an increase in refuge use). We reversed the sign of these values so that positive effect sizes would denote reductions in use of risky habitats, indicative of defence.

| Resource Response | Resource transition | Consumer transition | processes affected | Examples | Consumer Type | Potential costs to resource | Costs to consumer |
|---|---|---|---|---|---|---|---|
| Avoid Contact | ↑S-R | ↓ Q-A | Contact | burrow use when predators are present; vaccination from parasitic infection | All | reduced food intake and reproduction | Hunger |
| | ↓S-E | ↓ Q-A | Contact | camoflaged morphology of prey, leaving habitats with cues of parasite infestation | All | reduced food intake and reproduction | energy loss, hunger |
| Counter attack | ↑E-S | ↑A-Q | Attack failure<br>Resource Recovery S | Fleeing from approaching predators | Predator, micropredator | energy loss or allocation trade-offs | energy loss, hunger |
| | ↑E-S | ↑A death | Consumer death<br>Resource recovery to S | Deflecting parasitic propagules or micropredators; venemous biting of predators | All | energy loss or allocation trade-offs | Death |
| | ↑E-R | ↑A-Q | Attack failure<br>Resource recovery to S | fleeing approaching predators via dispersal to areas outside of consumer distribution | Predator | energy loss or allocation trade-offs | Energy loss, hunger |
| | ↑E-R | ↑A death | Consumer death<br>Resource recovery to S | Developing antibodies after deflecting parasitic propagules | All | energy loss or allocation trade-offs | Death, hunger |
| Combat consumption | ↑I-S | ↑C-Q | Consumption ended<br>Resource recovery to S | swatting, jolted body movement, posionous flesh | Micropredator | energy loss or allocation trade-offs | Failure, |
| | ↑I-S | ↑C death | Consumer Death<br>Resource recovery to S | Grooming, immune activation, habitat switching | Micropredator, parasite | energy loss or allocation trade-offs, reduced food intake and reproduction | Death |
| | ↑I-R | ↑C-Q | Consumption ended<br>Resource recovery to S | dispersal to areas outside of consumer distribution | Micropredator | energy loss or allocation trade-offs | Failure, hunger |
| | ↑I-R | ↑C death | consumer death<br>Resource recovery to S | Sterile immunity | Pathogens | energy loss or allocation trade-offs | Death, hunger |

**Table S3. Trait responses and their consequences for consumer-resource dynamics.** The three types of trait responses that predators and parasites may elicit (Column 1) have distinct effects on interaction dynamics (columns 2-3), in part because each response affects specific biological processes (Column 3). For columns 2 and 3: S = susceptible, E = exposed, I = ingested, R = resistant, Q = questing, A = attacking, C = consuming, as per Lafferty et al. 2015.  Arrows denote when responses increase transition rates and down arrows denote when trait responses decrease transition rates.

| Dataset | Factor/level | df | χ2 | d | se | t | p | CI lower | CI upper |
|---|---|---|---|---|---|---|---|---|---|
| **Full** | **interaction type** | 6 | 16.28 | | | | **0.003** | | |
| | predator-prey | | | 0.78 | 0.19 | 4.21 | **<.001** | 0.42 | 1.15 |
| | parasite-host | | | 0.21 | 0.19 | 1.11 | 0.265 | -0.16 | 0.57 |
| | combined | | | 0.89 | 0.21 | 4.32 | **<.001** | 0.49 | 1.29 |
| **Predators + Parasites** | **analysis scale** | 7 | 10.43 | | | | **0.015** | | |
| | group (predator) | | | 1.22 | 0.25 | 4.97 | **<.001** | 0.74 | 1.70 |
| | group (parasite) | | | 0.12 | 0.24 | 0.51 | 0.609 | -0.35 | 0.59 |
| | group (combined) | | | 1.44 | 0.28 | 5.22 | **<.001** | 0.90 | 1.98 |
| | individual (predator) | | | 0.42 | 0.22 | 1.93 | **0.054** | -0.01 | 0.85 |
| | individual (parasite) | | | 0.32 | 0.22 | 1.44 | 0.150 | -0.11 | 0.74 |
| | individual (combined) | | | 0.45 | 0.24 | 1.83 | 0.067 | -0.03 | 0.92 |
| | **trait type** | 6 | 8.84 | | | | 0.065 | | |
| | activity (predator | | | 1.11 | 0.23 | 4.89 | **<.001** | 0.67 | 1.55 |
| | activity (parasite) | | | 0.07 | 0.22 | 0.34 | 0.736 | -0.36 | 0.51 |
| | activity (combined) | | | 1.12 | 0.25 | 4.46 | **<.001** | 0.63 | 1.62 |
| | space use (predator | | | 0.33 | 0.37 | 0.90 | 0.369 | -0.40 | 1.06 |
| | space use (parasite) | | | 0.46 | 0.39 | 1.17 | 0.243 | -0.31 | 1.22 |
| | space use (combined) | | | 0.23 | 0.46 | 0.49 | 0.622 | -0.67 | 1.12 |
| | morphology/physiology (predator | | | 0.22 | 0.39 | 0.55 | 0.581 | -0.55 | 0.98 |
| | morphology/physiology (parasite) | | | 0.39 | 0.39 | 1.01 | 0.315 | -0.37 | 1.14 |
| | morphology/physiology (combined) | | | 0.73 | 0.40 | 1.84 | 0.066 | -0.05 | 1.50 |
| **Questing states** | **Interaction type** | 3 | 3.93 | | | | 0.140 | | |
| | predator-prey | | | 0.76 | 0.15 | 5.04 | **<.001** | 0.46 | 1.05 |
| | parasite-host | | | 0.12 | 0.29 | 0.42 | 0.677 | -0.45 | 0.70 |
| | combined | | | 0.53 | 0.50 | 1.06 | 0.291 | -0.45 | 1.51 |
| **Parasites only** | **Consumer state** | 4 | 8.43 | | | | **0.015** | | |
| | questing | | | 0.01 | 0.28 | 0.05 | 0.962 | -0.53 | 0.56 |
| | attacking | | | -0.28 | 0.30 | -0.96 | 0.339 | -0.86 | 0.30 |
| | consuming | | | 0.35 | 0.18 | 1.98 | **0.047** | 0.00 | 0.69 |
| | **Parasite strategy** | 5 | 1.32 | | | | 0.251 | | |
| | trophically-transmitted | | | 0.15 | 0.20 | 0.73 | 0.465 | -0.25 | 0.54 |
| | pathogen | | | 0.21 | 0.36 | 0.59 | 0.553 | -0.49 | 0.91 |

**Table S4. Estimated response magnitudes and the factors influencing responses.** Results of generalized linear mixed models used estimate the strength of non-lethal responses to predators and parasites and to explore factors influencing responses. Both the results from likelihood ratio tests testing the influence of factors (df, $\chi2$, p) and estimated effect sizes across different factor levels (d, se, t, p) are reported. The first set of models used the full dataset to test for differences in effect sizes for non-lethal responses to predators, parasites, or their combined presence. Models testing the influence of trait type and analysis scale omitted the combined treatments. The third set of model considered only data on responses to questing predators and parasites, and the fourth set examined factors influencing the strength of responses to parasites alone. Factors and factor levels with significant effects (denoted by $p < 0.05$) are highlighted in bold. df = degrees of freedom, d = estimated average effect (Hedge's d), se = standard error.

| Factor/level | df | $\chi2$ | p |
|---|---|---|---|
| interaction type:analysis scale | 14 | 5.31 | 0.070 |
| interaction type:trait type | 14 | 6.44 | 0.170 |

**Table S5.** Results of likelihood ratio tests (anova comparing full model to a model omitting the interaction) for the influence of interactions between explanatory variables in GLMMs using the full dataset (except combined treatments). The models used Hedge's d as the response and a Gaussian error structure since responses were normally distributed. Fixed effects in the full model included analysis scale, interaction type, and trait type, and we included interactions between interaction type and the other fixed effects in the full model. Interaction type denotes whether responses were made to predators or parasites. Analysis scale denotes whether responses were measured at the individual or group level. Trait type denotes the type of trait measured when estimating responses (activity, space use, morphology/physiology).

| **model** | **AICc** | **Δ AICc** | **weights** |
|---|---|---|---|
| d ~ interaction type + analysis scale | 312.21 | - | 0.52 |
| d ~ interaction type | 314.37 | 2.16 | 0.18 |
| d ~ interaction type + analysis scale + trait type | 314.71 | 2.50 | 0.15 |
| d ~ interaction type + trait type | 316.10 | 3.88 | 0.07 |
| d ~ analysis scale | 316.98 | 4.76 | 0.05 |
| d ~ 1 | 319.30 | 7.09 | 0.01 |
| d ~ analysis scale + trait type | 319.66 | 7.44 | 0.01 |
| d ~ trait type | 321.10 | 8.88 | 0.01 |

**Table S6.** Model ranking by AICc. Models of the strength of trait responses are shown in order of their performance, according to Akaike's Information Criterion for small sample sizes (AICc). Δ AICc = the difference in the AICc value from the top performing model.

| model | AICc | Δ AICc | weights |
|---|---|---|---|
| d ~ 1 + interaction type + analysis scale + trait type | 320.17 | - | 0.61 |
| d ~ 1 + interaction type + analysis scale | 322.34 | 2.17 | 0.20 |
| d ~ 1 + interaction type + trait type | 323.21 | 3.04 | 0.13 |
| d ~ 1 + interaction type | 326.65 | 6.48 | 0.02 |
| d ~ 1 + analysis scale + trait type | 327.26 | 7.09 | 0.02 |
| d ~ 1 + analysis scale | 328.19 | 8.02 | 0.01 |
| d ~ 1 + trait type | 330.08 | 9.91 | 0.00 |
| d ~ 1 | 331.82 | 11.65 | 0.00 |

**Table S6.** A list of candidate models for trait response magnitudes of larval amphibians, ranked according to their AICc value. Models used combinations of the following factors as fixed effects: interaction type (predator-prey, parasite-host, combined), analysis scale (individual, group), and trait type (activity, space use, morphological/physiological). Model ranking was done using the *glmulti()* function in the *glmulti* package in R.
